# Effects of childhood maltreatment on social cognition and brain functional connectivity in borderline personality disorder patients

**DOI:** 10.1101/461087

**Authors:** Xochitl Duque, Ruth Alcalá-Lozano, Jorge J. González-Olvera, Eduardo A. Garza-Villarreal, Francisco Pellicer

## Abstract

Borderline personality disorder (BPD) is a chronic condition characterized by high levels of impulsivity, affective instability, and difficulty to establish and manage interpersonal relationships. This paper assessed differences in performance on social cognitive paradigms (MASC, RMTE) and how it related to child abuse. Specifically, it evaluated the relationship between performance on cognitive paradigms and baseline brain connectivity in patients with BPD, compared to healthy controls.

BPD patients had higher levels of childhood maltreatment, increased impulsivity and aggression, and more dissociative symptoms than control subjects. For the sexual abuse subdimension, there were no differences between the BPD and the control groups, but there was a negative correlation between MASC scores and total childhood maltreatment levels, as well as between physical abuse, physical negligence, and MASC. Both groups showed that the higher the level of childhood maltreatment, the lower the performance on the MASC social cognitive test. Further, in the BPD group, there was hypoconnectivity between the structures responsible for emotion regulation and social cognitive responses that have been described as part of the frontolimbic circuitry. The more serious the child abuse, the lower the connectivity.

## 1 Introduction

Borderline personality disorder (BPD) is a chronic psychiatric condition characterized by high levels of impulsivity and affective instability, as well as a marked difficulty to establish and manage interpersonal relationships (1,2). In patients with this multifactorial disorder, a genetic vulnerability has been identified (1,2). This vulnerability may interact with environmental factors such as lower quality parental care (3,4) and a history of child abuse, which is present in a large number of research subjects with BPD and has been proposed to be proposed to be a contributing factor of the disorder(5).

Chronic difficulties in interpersonal relationships are a BPD characteristic and have been studied within the social cognition construct (6–8). With regard to performance in recognizing the emotions of others, some studies have found higher levels of performance on tests among BPD patients, including the reading the mind in the eyes test (RMTE) (9,10), while other studies have not reported any differences in the ability to infer the mental states of self and others, compared to controls (11–13). Still, other authors that have used ecological paradigms such as the Movie for the Assessment of Social Cognition (MASC) have found a deficit in social cognition (7)

The neurobiological substrate of social cognition in BPD has been studied by task-related neuroimaging studies such as the RMET paradigm and stimuli adaptations that test Theory of Mind (ToM). These studies showed BPD patients have lower activation in areas within the temporal lobe, the superior and medial frontal regions, the cingulate cortex, parietal cortex, hippocampus, and the insula, as well as higher activation in bilateral amygdala, left temporal pole, medial frontal gyrus, right middle and superior temporal gyrus, left precuneus, left middle occipital gyrus and right insula compared to controls (10,14,15). In addition, a lower brain response has been reported in the BPD group in the left superior temporal sulcus and gyrus in response to the modified version of the Multifaceted Empathy Test (MET) (16). The functional connectivity describes the neuronal activity correlation between different brain regions. Most studies describe correlations observed between low-frequency fluctuations (<0.1 Hz) at basal state, which are organized in intrinsic neural networks which are the same previously described in task-related research (17–19). One of these networks is the default mode network (DMN), which shows a decreased connectivity in the precuneus, (14) and the right posterior cingulate (20), as well as hyperconnectivity in the medial prefrontal cortex, the anterior cingulate cortex, and the posterior precuneus/cingulate in BPD compare to healthy subjects (21).

Most regions where differences were found in the brain function in BPD form part of the frontolimbic circuit. Dysfunction of frontolimbic circuitry is one of the most accepted models to explain the BPD symptoms, including emotional dysregulation and social cognition deficits (22). This same circuit has been related to morphologic and functional brain changes associated with a history of child abuse (23). Previous research showed gray matter volume reduction in orbitofrontal cortex and temporal regions (24,25) and hyperactivation in response to affect-laden stimuli in the right amygdala (26) related to childhood maltreatment as well as an effect in functional connectivity (27,28).

Even though brain activation has been studied regarding social cognition tasks, the relationship between functional connectivity at resting state and its association with the performance in such tasks has not been explored. The inclusion of the childhood maltreatment variable may offer information that could contribute to understanding the heterogeneity of clinical and neuroimaging results in BPD studies (29). The primary goal of this paper was to assess differences compared to healthy controls in the clinical performance of social cognitive paradigms and functional connectivity in resting state and how it related to child maltreatment levels.

## 2. Materials and Methods

For our study, we included 18 patients diagnosed with BPD and 15 controls without any psychiatric diagnosis (CN) in a cross-sectional design. Both groups were matched by age and education. Due to the higher prevalence of the psychiatric diagnosis among women, all study participants were women (30) and right-handed. Participants were recruited from the outpatient clinic of the Institute for Social Security and Services for State Workers (ISSSTE). We also recruited 4 participants from Instituto Nacional de Psiquiatría “Ramón de la Fuente Muñiz” from an ongoing study (31). The protocol was approved by the Ethics Committee of the ISSSTE (317.2017_P_2017) and the Ethics Committee of the Instituto Nacional de Psiquiatría “Ramón de la Fuente Muñiz”. All the participants signed an informed consent form, and the study followed the guidelines in the Declaration of Helsinki.

Patients diagnosed with BPD between 18 and 45 years old were included. The BPD diagnosis was established by the attending psychiatrist and corroborated by a psychiatrist with experience in BPD, who used the Diagnostic Interview for Borderline Revised (cut-off of 6)(32). To determine comorbidity, we used the Spanish version of the Mini International Neuropsychiatric Interview (MINI)(33). To obtain a representative sample of the clinical population, the study included patients with Major depressive disorder (MDD), posttraumatic stress disorder comorbidity (PTSD) and the use of medication. Exclusion criteria were disorder caused by use of addictive substances in the last six months, bipolar disorder diagnosis, schizophrenia, obsessive-compulsive disorder, eating disorders, and mental disability as described by the attending physician. For the control group, psychopathologies were ruled out with the MINI. Diagnosis of Axis II disorders was ruled out by means of SCID-II screening, and positives were evaluated by the psychiatrist.

We measured the social cognition construct using the Spanish version(34) of Movie for the Assessment of Social Cognition (MASC)(35). This version is a 16-minute video depicting social situations where the protagonists’ emotions, thoughts, and social intentions are assessed through 46 multiple-choice questions. For each question, there is only one right answer. Mistakes were classified as hyper-, hypo-, and lack of mentalization. The test has high inter-rater (ICC = 0.99) and test-retest reliability (r = 0.97) and is highly consistent among observers (Cronbach’s α = 0.86) (35). The video was provided by the author of the Spanish version (Guillermo Lahera; Universidad de Alcalá, Madrid, Spain) and professionally dubbed into Mexican Spanish with an adaptation to the Mexican accent and words. In addition, the study used the Reading the Mind in the Eyes test (RMTE) to assess the ability to infer mental states with information from the eye gaze in pictures (36). Each participant was asked to choose one of four descriptions of mental states for each picture. The Barratt Impulsivity Scale (BIS-11), Buss-Perry Aggression Questionnaire (BPAQ) and Dissociative Experiences Scale (DES) were applied. To determine whether there was a history of childhood trauma, the Spanish version of the Childhood Trauma (self-administered) Questionnaire (CTQ) (37) was used.

### 2.1 Magnetic resonance imaging

Imaging data were obtained using a Phillips Ingenia 3 T with a 32-channel phased-array head coil. We acquired structural and resting state functional (fMRI) sequences. For the resting-state fMRI (rsfMRI), participants were instructed to remain quiet, keep their eyes open, without thinking of anything in particular and were presented with a white cross on a black background. T2*-weighted echo planar images were acquired with the following parameters: 36 axial slices, repetition time = 2000 ms, echo time = 30 ms, flip angle = 75°, field of view = 240 mm, slice thickness = 3.0 mm, acquisition matrix = 80 × 80, and voxel size = 3.0 × 3.0 × 3.0 mm^3^. Structural T1-weighted images were acquired with a repetition time = 7 ms, echo time = 3.5 ms, flip angle = 8°, field of view = 240 mm, slice thickness = 1.0 mm, acquisition matrix = 240 × 240, and voxel size = 1.0 × 1.0 × 1.0 mm^3^.

### 2.2 Statistical analysis of clinical measures

The statistical software SPSS-X version 22.0 for Windows, PC, was used for the analyses. We visually inspected the clinical data and used the Shapiro-Wilks test to assess for normality. We first compared the scores from the social cognition variables (MASC and RMTE), CTQ, BPAQ, BIS-11 and DES scale between the BPD and CN groups using a paired t-test (Mann-Whitney U test for non-normal variables) with alpha of 0.05. We then performed Pearson’s correlation between the MASC and RMTE and CTQ scores to search for a possible relationship between childhood maltreatment and social cognition. Finally, we created a new nominal variable using the MINI with the following factors: BPD with depression (n = 11), BPD without depression (n = 7), and CN. Then we used a one-way ANOVA to find differences in social cognition variables between the groups.

### 2.3 Resting state functional connectivity preprocessing and analysis

Data were preprocessed and analyzed using the CONN-fMRI Functional Connectivity toolbox (38). The preprocessing pipeline prior to the analysis included: functional realignment and unwrap (subject motion estimation and correction, functional center to (0,0,0) coordinates (translation), slice-timing correction, detection of motion artifact sources with ART (Artifact Detection Tools; developed by Stanford Medicine, Center for Interdisciplinary Brain Sciences Research) (Time points exceeding the movement threshold of 2 mm or a global signal Z-value of 9 were defined as outliners), direct segmentation and normalization (simultaneous Gray/White/CSF segmentation and normalization to MNI space), and smoothing (5-mm FWHM Gaussian filter). With a general linear model, nuisance variables were regressed out. The nuisance variables included were: subject motion parameters, raw white matter, and cerebrospinal fluid signals. To correct for physiological noise, we used the CompCor method (39). Signal time series were band-pass filtered between 0.008 and 0.09 Hz.

To assess baseline functional connectivity (rs-FC), we carried out a seed-based correlation analysis. The seed regions were defined in CONN using an 8 mm kernel sphere (Figure 1). The definition of the seeds was based on previous BPD results and regions associated with mentalization, especially those in the DMN (10,14,15,21,40,41) (For details see Supplementary Material Table 1S). The whole-brain individual correlation maps were computed with the average value of the BOLD signal time course in resting state in each seed region, and correlation coefficients were estimated with the BOLD signal time course for each voxel. A normal distribution of the resulting coefficients was obtained with the Fisher transformation, and correlation maps (functional connectivity) were obtained for each seed region and subject. The correlation maps for each seed were used to carry out a second-level between-groups contrast GLM using age as a covariate. All contrasts were corrected for multiple comparisons with the false discovery rate, with a p-threshold of 0.05 for each test and cluster. Finally, we extracted the Z-maps (Fisher-transformed connectivity values) for each significant cluster in each subject to perform Pearson correlation between functional connectivity and clinical measures. Previous research indicated a higher likelihood of false positives resulting from multiple comparisons. This was particularly the case of studies that correlated brain activation with behavioral variable results. Thus, the study corrected for multiple comparisons (42,43).

**Figure 1.**
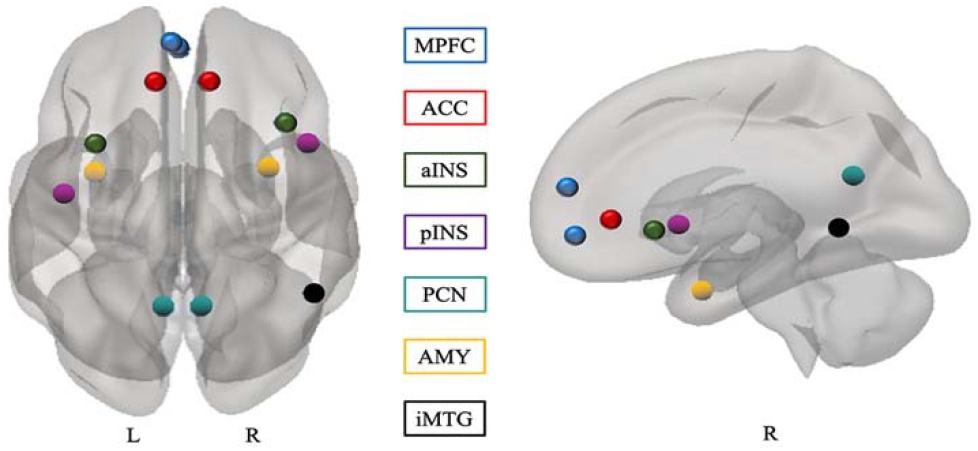
Seeds used in this study. This figure shows the seeds; all seeds were defined based on previous BPD studies (see Table S1 in the Supplementary Materials). MPFC, medial prefrontal cortex; ACC, anterior cingulate cortex; PCN, Precuneus; iMTG, inferior middle temporal gyrus; AMY, amygdala; INS, insular cortex; L, left; R, right; a, anterior; p, posterior

## 3. Results

### 3.1 Clinical measures

The psychiatric comorbidity and medications of the BPD group are summarized in Supplementary Material Table 2S. Compared to the controls, BPD patients showed an increase in impulsivity, aggression levels, and dissociative symptoms, and higher scores on the CTQ. Regarding abuse subdimensions, there were no significant differences in sexual abuse between the groups (Table 1). There was a negative correlation between MASC scores and total CTQ score; for the subdimensions, there was a negative correlation between physical abuse, physical negligence, and total MASC, as shown in Table 2.

**Table 1.** Demographic and clinical characteristics of BPD patients and healthy participants groups

|  | BPD (n= 18) |  | HC (n= 15) |  | p-value |
| --- | --- | --- | --- | --- | --- |
| Age in years | 31.17 | ± 9.52 | 32.80 | ±8.65 | p=.613 <sup>a</sup> |
| Years in education | 15.06 | ±2.226 | 15.29 | ±2.58 | p=.790 <sup>a</sup> |
| DES | 30.28 | ±19.93 | 9.94 | ±7.63 | p= .003 <sup>a</sup> |
| BPAQ | 91.25 | ±20.95 | 72.08 | ±19.85 | p= .021 <sup>a</sup> |
| BIS-11 | 63.50 | ±14.09 | 50.15 | ±15.50 | p= .022 <sup>a</sup> |
| CTQ TOTAL | 59.06 | 18.63 | 38.87 | 8.50 | p=.001 <sup>a</sup> |
| Emotional Abuse | 16.56 | 5.31 | 9.53 | 2.41 | p=.000 <sup>a</sup> |
| Emotional neglect | 11.44 | 3.63 | 7.93 | 3.15 | p=.006 <sup>a</sup> |
| Physical neglect | 9.50 | mdn | 6.00 | mdn | P=.006 <sup>b</sup> |
| Physical abuse | 7.00 | mdn | 9.00 | mdn | P=.059 <sup>b</sup> |
| Sexual abuse | 9.00 | mdn | 5.00 | mdn | p=.137 <sup>b</sup> |
| MASC total correct | 30.00 | ±5.3 | 31.03 | ±2.58 | p=.449 <sup>a</sup> |
| Overmentalizing errors | 6.00 | ±2.89 | 5.23 | ±2.7 | p=.538 <sup>a</sup> |
| “reduced ToM” errors | 4.0 | mdn | 6.0 | mdn | p=.686 <sup>b</sup> |
| “no ToM” errors | 3 | mdn | 3 | mdn | p=.714 <sup>b</sup> |
| RMTE | 25.06 | ±3.26 | 25.73 | ±4.3 | P=.614 <sup>a</sup> |

Data are presented in means ± standard deviation unless otherwise specified. BPD, borderline personality disorder; HC, healthy controls; mdn , median; DES, Dissociative Experiences Scale; BPAQ, Buss-Perry Aggression Questionnaire; BIS-11, Barratt impulsiveness scale; CTQ, Childhood Trauma Questionnaire; MASC, Movie for the Assessment of Social Cognition; ToM, theory of mind; RMTE, Reading the mind in the eyes.
<sup>a</sup> Two-sample two-tailed t-test.
<sup>b</sup> Mann–Whitney U test.

**Table 2.** Table of correlations between variables of social cognition and CTQ scores.

|  | 1 | 2 | 3 | 4 | 5 | 6 | 7 | 8 | 9 | 10 | 11 |
| --- | --- | --- | --- | --- | --- | --- | --- | --- | --- | --- | --- |
| 1 MASC total correct | 1 |  |  |  |  |  |  |  |  |  |  |
| 2 MASC overmentalizing errors | -.02 | 1 |  |  |  |  |  |  |  |  |  |
| 3 MASC “reduced ToM” errors | -.77<sup>**/++</sup> | -.43<sup>*/++</sup> | 1 |  |  |  |  |  |  |  |  |
| 4 MASC “no ToM” errors | -.50<sup>**/++</sup> | -.40<sup>*/+</sup> | .48<sup>**/++</sup> | 1 |  |  |  |  |  |  |  |
| 5 RMTE | .35<sup>*/+</sup> | .37<sup>*/+</sup> | -.42<sup>*/++</sup> | -.20 | 1 |  |  |  |  |  |  |
| 6 CTQ_Total | -.38<sup>*/+</sup> | -.04 | .34<sup>*/+</sup> | .09 | -.15 | 1 |  |  |  |  |  |
| 7 Physical neglect | -.38<sup>*/+</sup> | -.05 | .42<sup>*/++</sup> | .03 | -.23 | .75<sup>**/++</sup> | 1 |  |  |  |  |
| 8 Emotional Abuse | -.30 | .13 | .18 | -.04 | -.15 | .86<sup>**/++</sup> | .59<sup>**/++</sup> | 1 |  |  |  |
| 9 Emotional neglect | -.24 | -.34<sup>*/+</sup> | .36<sup>*/+</sup> | .19 | -.29 | .69<sup>**/++</sup> | .62<sup>**/++</sup> | .56<sup>**/++</sup> | 1 |  |  |
| 10 Physical abuse | -.40<sup>*/+</sup> | -.06 | .37<sup>*/+</sup> | .16 | -.04 | .88<sup>**/++</sup> | .59<sup>**/++</sup> | .74<sup>**/++</sup> | .47<sup>**/++</sup> | 1 |  |
| 11 Sexual abuse | -.18 | .04 | .12 | .05 | .02 | .69<sup>**/++</sup> | .31 | .40<sup>**/+</sup> | .22 | .57<sup>**/++</sup> | 1 |
| Mean | 30.53 | 5.67 | 5.97 | 3.39 | 25.73 | 49.55 | 7.79 | 13.24 | 9.82 | 9.30 | 9.39 |
| Standard Deviation (SD) | 4.31 | 2.74 | 3.59 | 2.63 | 3.38 | 17.99 | 3.18 | 5.56 | 3.81 | 4.92 | 5.49 |

$n = 33$ ; \*\*: $p < 0,01$ (bilateral); \*: $p < 0,05$ (bilateral); +: FDR < 0.1; ++: FDR < 0.05; MASC, Movie for the Assessment of Social Cognition; ToM, theory of mind; RMTE, Reading the mind in the eyes; CTQ, Childhood Trauma Questionnaire.

In the BPD group, depression manifested in different ways. Depressed BPD subjects had lower performance on the MASC (M= 27.73, SD = 5.350) and a decrease in the mean of −5.84, 95% CI [−11.02, −.67](p = 0.026), compared to non-depressed BPD subjects (M= 33.57, SD = 3.15), who even performed better than the controls on the MASC (M= 31, 17, SD = 2.57); the difference was, however, not statistically significant (p = 0.53), as determined by one-way ANOVA for the three groups, F (2, 14.041) = 4.09, p < 0.040. No significant differences in RMTE scoring were found between the groups, F (2, 30) = .479, p = 0.305.

### 3.2. Functional connectivity

Hypoconnectivity was found between limbic regions that play a role in emotional and affective regulation and social responses in BPD patients. A hyperconnectivity was observed between the medial prefrontal cortex and the left superior parietal lobe (Table 3 and Figure 2). There were no statistically significant differences in the connectivity values between non-depressed and depressed BPD subjects (Supplementary Material Table 2S).

**Figure 2.**
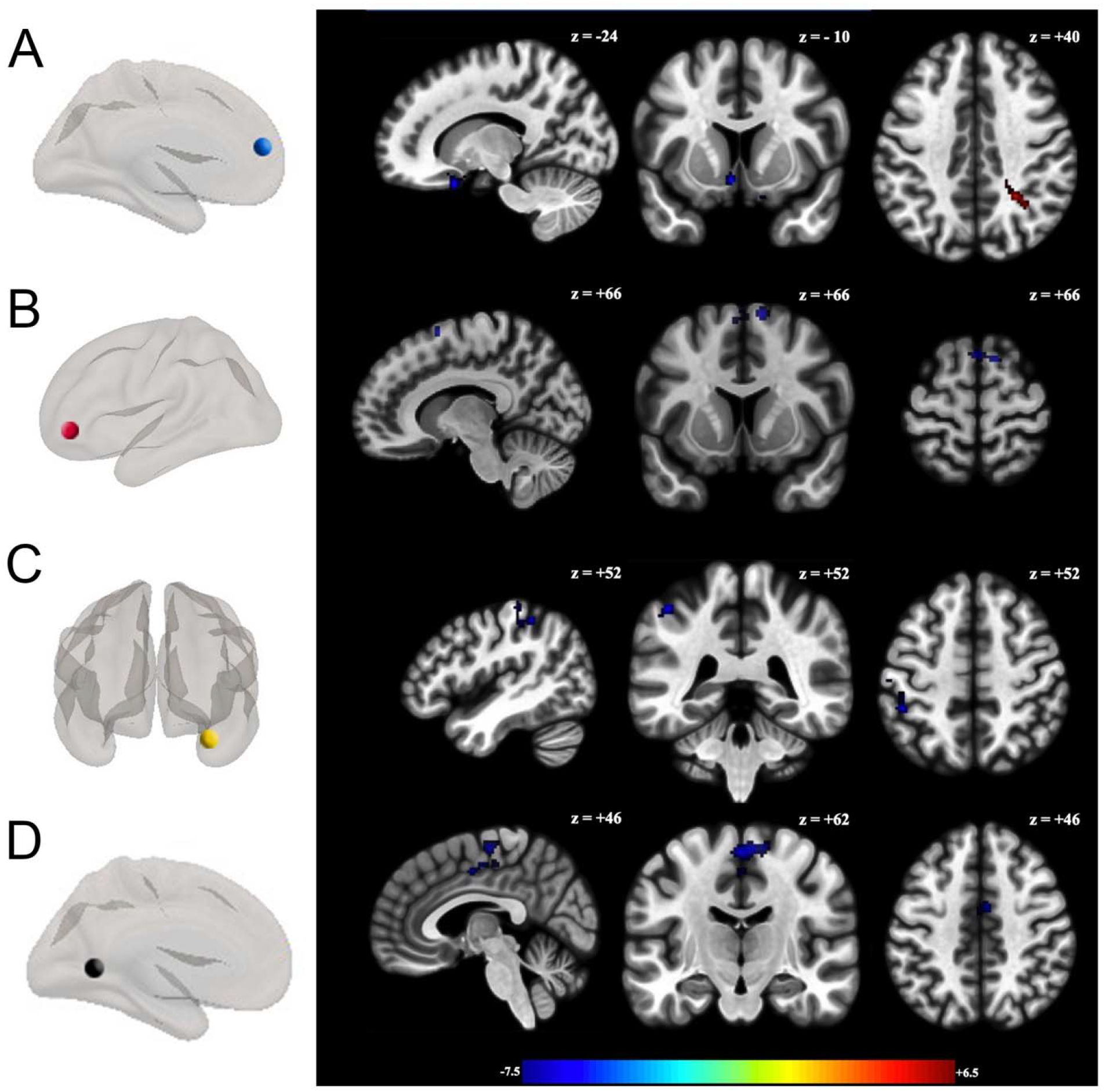
Seeds showing significant functional connectivity differences between groups with the (A) left medial prefrontal cortex, (B) anterior cingulate cortex right; (C) right amygdala; and (D) inferior Middle temporal gyrus between BPD patients and healthy controls controlling for age. All analyzed contrasts where corrected by multiple comparisons using the false discovery rate (FDR) at 0.05. Blue and Orange/hot represent decreased and increased functional connectivity, respectively. The color bar indicates the *t*-value. Details of the clusters are shown in Table 3

**Table 3.** Seeds showing significant functional connectivity differences between groups (BPD> CN) controlling for age

| Seed | Regions | Peak voxel coordinate | | | Cluster size | $\beta$ | t-scores | Cluster significance (FDR-corrected, threshold of $p=0.05$) | Connectivity |
| --- | --- | --- | --- | --- | --- | --- | --- | --- | --- |
|  |  | x | y | z |  |  |  |  |  |
| MPFC_L | SPL_R | +26 | -48 | +40 | 61 | 0.21 | 6.05 | 0.0383 | higher |
|  | NaC.L, SubCalC, P_L, Cd_L, OFC.L | -06 | +10 | -10 | 80 | -0.17 | -5.82 | 0.0213 | lower |
|  | OFC.R, SubCalC | +14 | +16 | -24 | 76 | -0.19 | -7.34 | 0.0213 | lower |
| ACC_R | SFG_L, SFG_R | +10 | +10 | +66 | 110 | -0.18 | -5.42 | 0.0059 | lower |
| AMYG-R | SI_L, aSMG_L, SPG_L pSMG_L | -46 | -40 | +52 | 154 | -0.16 | -5.72 | 0.0010 | lower |
| iMTG-R | M1_R, M1_L, SMA_R, SMA_L | +00 | -16 | +62 | 180 | -0.23 | -4.62 | 0.0008 | lower |
|  | SMA_R, M1_L, M1_L, M1_R, ACC | +04 | -04 | +46 | 131 | -0.23 | -4.66 | 0.0034 | lower |

$\beta$ , effect size (positive effects represent higher connectivity; negative effects represent lower connectivity); T, T-value; p-FDR, p corrected false discovery rate; L, left; R, right; a, anterior; p, posterior; i, inferior; s, superior; **Seeds**: MPFC, medial pre-frontal cortex; ACC, anterior cingulate cortex; AMYG, amygdala; MTG Middle temporal gyrus. **Correlated areas**: Cd, Caudate; M1, precentral gyrus (primary motor cortex); NaC, Accumbens; OFC, Orbitofrontal cortex; P, Putamen; SI, Postcentral gyrus (primary somatosensory cortex); SFG, superior frontal gyrus; SubCalC, subcallosal cortex; SMG, supramarginal gyrus; SPL, superior parietal lobe; SMA, supplementary motor area. All analyzed contrasts were corrected by multiple comparisons using the false discovery rate (FDR) at 0.05.

### 3.3 Correlation between clinical and functional connectivity

We used Fisher-transformed connectivity values of the seven clusters identified with the comparative analysis of the groups and correlated with the MASC, Movie for the Assessment of Social Cognition (MASC), Reading the mind in the eyes (RMTE) and Childhood Trauma Questionnaire (CTQ) scores. For the largest number of regions studied, a negative correlation was found between functional connectivity and the total levels of child abuse, as well as some subdivisions of abuse as shown in Table 4. That is, higher levels of child abuse are related to less connectivity in these regions (Figure 3).

**Figure 3.**
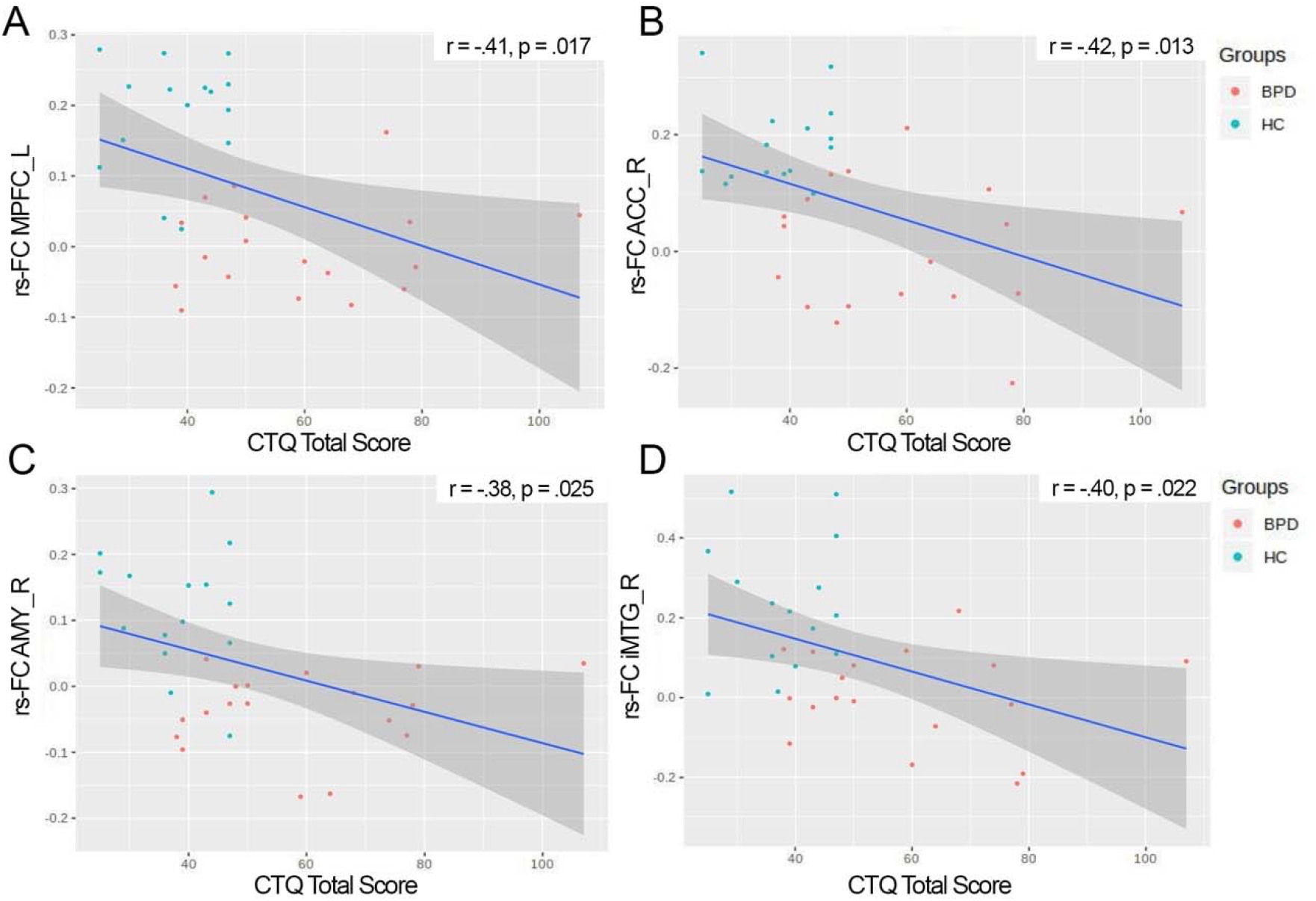
Correlation between childhood trauma (total CTQ score) and functional connectivity. The figure shows connectivity values for seeds and cluster: (A) MPFC_L, left medial prefrontal cortex, cluster (x = +14, y = +16, z = −24); (B) ACC_R, right anterior cingulate cortex, cluster (x = +10, y = +10, z = 66) : (C) AMY_R, right amygdala, cluster (x = −46, y = −40, z = +52) and (D) iMTG_R, inferior Middle temporal gyrus (x = +00, y = −16, z = +62). rs- = correlation coefficient. BPD = borderline personality disorder, HC = healthy control.

**Table 4.** Correlations between functional connectivity and clinical measures

| Cluster<br>MNI | MPFC_L<br>+26 -48, +40 | MPFC_L<br>-06 +10 -10 | MPFC_L<br>+14 +16 -24 | ACC_R<br>+10 +10 +66 | AMYG-R<br>-46 -40 +52 | iMTG-R<br>+00 -16 +62 | iMTG-R<br>+04 -04 +46 |
| --- | --- | --- | --- | --- | --- | --- | --- |
| Clinical Measures | r (p value) | r (p value) | r (p value) | r (p value) | r (p value) | r (p value) | r (p value) |
| MASC total correct | -0.135 (0.452) | 0.220 (0.218) | 0.201 (0.261) | 0.355 (0.042) | -0.050 (0.779) | 0.095 (0.597) | 0.1263 (0.483) |
| Overmentalizing errors | 0.011 (0.951) | -0.146 (0.414) | -0.139 (0.437) | -0.248 (0.164) | -0.210 (0.240) | -0.092 (0.607) | -0.048 (0.790) |
| “reduced ToM” errors | 0.062 (0.729) | -0.068 (0.707) | -0.090 (0.617) | -0.130 (0.470) | 0.139 (0.438) | 0.030 (0.864) | 0.030 (0.865) |
| “no ToM” errors | -0.072 (0.688) | -0.143 (0.426) | -0.181 (0.312) | -0.081 (0.652) | 0.142 (0.430) | 0.032 (0.856) | -0.161 (0.370) |
| RMTE | -0.362 (0.038) | 0.212 (0.234) | 0.272 (0.124) | 0.242 (0.174) | 0.031 (0.861) | -0.078 (0.663) | -0.064 (0.721) |
| CTQ TOTAL | **0.443 (0.009)<sup>+</sup>** | -0.248 (0.162) | **-0.411 (0.017)<sup>+</sup>** | **-0.427 (0.013)<sup>+</sup>** | -0.389 (0.025) | **-0.409 (0.018)<sup>+</sup>** | **-0.395 (0.022)** |
| Emotional Abuse | **0.480 (0.004)<sup>+</sup>** | **-0.393 (0.023)** | **-0.431 (0.012)<sup>+</sup>** | **-0.503 (0.002)<sup>+</sup>** | **-0.495 (0.003)<sup>+</sup>** | **-0.508 (0.002)<sup>+</sup>** | **-0.483 (0.023)** |
| Emotional neglect | 0.309 (0.079) | -0.330 (0.060) | **-0.471 (0.005)<sup>+</sup>** | -0.231 (0.194) | -0.288 (0.102) | **-0.427 (0.013)<sup>+</sup>** | **-0.437 (0.004)<sup>+</sup>** |
| Physical neglect | **0.469 (0.005)<sup>+</sup>** | -0.323 (0.066) | **-0.426 (0.013)<sup>+</sup>** | -0.340 (0.052) | -0.291 (0.099) | -0.236 (0.186) | -0.176 (0.326) |
| Physical abuse | 0.234 (0.188) | -0.142 (0.430) | -0.233 (0.191) | -0.335 (0.056) | -0.227 (0.203) | -0.211 (0.237) | -0.258 (0.145) |
| Sexual abuse | 0.267 (0.132) | 0.127 (0.480) | -0.127 (0.480) | -0.232 (0.193) | -0.199 (0.266) | -0.203 (0.256) | -0.168 (0.349) |

MPFC, medial pre-frontal cortex; ACC, anterior cingulate cortex; AMYG, amygdala; MTG Middle temporal gyrus; Numbers represents: Pearson coefficient, p values; MASC, Movie for the Assessment of Social Cognition; ToM, theory of mind; RMTE, Reading the mind in the eyes; CTQ, Childhood Trauma Questionnaire. +: FDR < 0.1; No significant values were observed after FDR.05

## 4. Discussion

We found that as the level of childhood maltreatment increased, the performance on the MASC social cognitive test decreased in both the BPD and the control groups. In addition, there was hypoconnectivity between structures associated with emotion regulation and social cognitive responses in the BPD group. Connectivity decreased as levels of child abuse increased.

Our findings relate to another study, where they showed that child abuse has an impact on the skills necessary to develop stable and long-lasting interpersonal relationships(44). It has been well established that the development of social cognition is linked to that of emotional and affective communication through primary caregivers and an environment that is safe and free from excessive stress-conditions that do not exist in the case of child abuse (45). Several studies have documented that a disruption in the relationships between children and primary figures or extremely stressful environments activate the hypothalamic-hypophyseal axis by releasing and activating several mechanisms that have an effect on the brain (46,47). This physiological environment of extreme stress interferes with the integration of mental representations during development, thus disrupting the concepts of self and other and producing an unrealistic, unstable, and disproportionate representation of the affection perceived and expressed (3). These circumstances arise in relational contexts, a fact that accounts for BPD patients’ clinical characteristics. These traits, in our study, may have been expressed as changes in connectivity, aggression, impulsivity, and dissociative symptoms, as compared to the control group.

We did not find differences between social cognitive tasks for both paradigms, which contrasts with the ongoing controversy regarding a BPD patient’s ability to read mental states (10,48). Data collected with the MASC goes beyond the underlying process of recognizing emotion in social interactions with information from the eye gaze that RMTE evaluates (36). It also includes an assessment of the content of the mental state of the “other,” based on contextual information and elements that are not physically evident (49). This suggests that the instrument is ideal as it reflects real-life situations. Nevertheless, the two paradigms evaluate the cognitive dimension of the social cognitive process, and it is impossible to rule out the limitations of the affective dimension that are associated with emotional regulation and the difficulties in distinguishing between self and the other in BPD subjects (50). Our study showed that depression is associated with decreased social cognitive performance. Although other research has found similar results (51), some authors have not found differences between the BPD and the control groups and have surmised that rather than a state, social cognition performance is a trait (52).

### 4.1 Functional Connectivity Results

Our results identified differences in the organization of brain activation patterns between the groups, mainly hypoconnectivity between the regions explored in BPD. These regions are related to a broad range of cognitive and emotional processes most of which play a role in social cognition. Besides, the regions that we studied are a part or are related to the default mode network, which is activated by processes that involve thought forms created by self as autobiographical memory, planning for the future, and inferring one’s mental states and those of “others” (53). Activity in the medial prefrontal cortex (MPFC) is associated mainly with the ability to differentiate between self and the other, the detection of one’s own emotional state and the ability to mentalize (53,54). The temporal lobe has been implicated in the processing of language and facial expressions, and it plays an essential role in the process of inference of mental states (55,56). The anterior cingulate cortex (ACC) participates in tasks of behavior monitoring, detection and prediction of error, decision making and processes related to self-evaluation, especially in social contexts (57,58) in conjunction with the amygdala which is a critical structure for the emotional regulation and has been studied extensively in BPD (59–61). All these processes are essential for the success of interpersonal relations. The differences in the connectivity of these structures may explain the impairment in differentiating between self and others exhibited by the syndrome of identity diffusion that characterizes borderline personality organization and could underlie primitive defense mechanisms such as splitting (62).

Some of the results in this study agree with previous task-based studies. The decrease of activity in the temporal region can be found in previous studies(63,64). Regarding the amygdala, our findings are in accordance with previous research (65), but evidence has been inconsistent (66). On the other hand, studies have found a hyperactivation in MPFC and ACC (21,67) even though there are results that show a hypoactivation of these structures at resting state (68,69) and task-based studies (14,65). In this study, although there were differences in the brain connectivity in these regions with such an importance for the social behavior, we did not observe a correlation with the clinical variables of social cognition. The differences in the reading of the mental states in BPD are observed especially under the effects of emotional stress (70,71). In that sense, we assume that the clinical performance in social cognition tasks related to brain organization at rest could vary under intense emotional states. Stress is associated with an abnormal pattern of deactivation of intrinsic neural networks(72,73), which could be associated with variations in the performance of social skills. However, in this study, it was not possible to show the effect of stress on brain organization.

Finally, we found an effect of childhood abuse on brain functional connectivity. The higher the level of child abuse in the patients, the lower their brain connectivity. Although the effect of child maltreatment on brain structures has been widely documented (27,74,75), the mechanism with which child abuse might have an impact on organization and functional connectivity is still unclear. A possible explanation is that child abuse is a factor associated with brain remodeling rather than a harmful factor per se (29,76), especially with corticolimbic structures, as shown by a preclinical study of adolescent rats (77). This “modeling” effect on brain organization is present especially during critical stages and processes, such as pruning that is necessary for normal brain development (78). In the first two years of life, a synaptic overproduction occurs in the brain, followed by remodeling through pruning; these processes continue into adolescence (79). Although remodeling occurs due to cellular programming, research has reported that pruning in this second phase is highly sensitive to experience (80), including stress, because of the effect of inflammation mechanisms on glial cells (81,82). This is consistent with the new paradigm that regards the brain as an active system that self-organizes dynamically based on the information that it receives (83). This has been studied in schizophrenic and autistic subjects, for whom dysfunctional pruning has been proposed (79,84). Nonetheless, in BPD patients remodeling and differential pruning would be associated with stressful childhood events or a lack of proper parenting. This proposition helps to understand BPD as the result of maladaptive brain remodeling produced by the effect of traumatic experiences on brain development.

## 5. Conclusion

BPD patients endured more child abuse than the controls, which correlated with poorer performance on the MASC social cognitive test and lower connectivity between structures involved in emotion regulation and social cognitive responses, that are part of the frontolimbic circuitry. The rsfMRI results provide information about internal baseline processed that seem altered in BPD patients. The higher the level of child abuse in the patients, the lower their brain connectivity. We need to further study these results, and there is a need to find if the introduction of safeguards to avoid abuse and stress in such critical periods as are childhood and adolescence are beneficial, and patients may recover from these harmful effects.

## Acknowledgements

The study was supported by grants through the institutional program E015 and from the National Council of Science and Technology of Mexico CONACYT-FOSISS project N0. 289831, No. 0201493 and CONACYT-Cátedras project No. 2358948. We thank the people who support this project in one way or another: Ana R. Villaseñor, Diego Angeles Fiacro Jiménez Ponce. Finally, we thank the study participants for their time and patience

## Ethics Statement

This study was carried out in accordance with the recommendations of Ethics Committee, with written informed consent from all subjects. All subjects gave written informed consent in accordance with the Declaration of Helsinki. The protocol was approved by the Ethics Committee of both institutions the Instituto de Seguridad y Servicios Sociales de los Trabajadores del Estado y del Instituto Nacional de Psiquiatría “Ramón de la Fuente Muñiz”.

## Author Contributions

XD, FP, and JG-O were involved in the design of the research protocol. XD, RA, and EG-V contributed to acquisition and analysis of data; XD, FP, and EG-V drafted the manuscript, and all authors contributed revising and approved it for publication

